# Heterodimerization-dependent secretion of BMPs in *Drosophila*

**DOI:** 10.1101/2022.08.04.502599

**Authors:** Milena Bauer, Gustavo Aguilar, Kristi A. Wharton, Shinya Matsuda, Markus Affolter

## Abstract

Combinatorial signaling is key to instruct context-dependent cell behaviors. During embryonic development, adult homeostasis and disease, Bone Morphogenetic Proteins (BMPs) act as dimers to instruct specific cellular responses. BMP ligands can form both homo- or heterodimers; however, obtaining direct evidence of the endogenous localization and function of each form has proven challenging. Here, we make use of precise genome editing and direct protein manipulation via protein binders to dissect the existence and functional relevance of BMP homo- and heterodimers in the *Drosophila* wing imaginal disc. This approach revealed *in situ* the existence of Dpp (BMP2/4)/Gbb (BMP5/6/7/8) heterodimers. We found that Gbb, despite being produced by all the cells of the wing imaginal disc, is only secreted by the cells in which Dpp is expressed. Dpp and Gbb form a gradient of heterodimers whereas neither Dpp nor Gbb homodimers are evident under endogenous physiological conditions. We find that the formation of heterodimers is critical for obtaining optimal signaling and long-range BMP distribution in the developing wing. These results indicate that Dpp/Gbb heterodimers are the active signal required for epithelial patterning and growth.

## Introduction

Bone morphogenetic proteins (BMPs) represent an ancient group of signaling ligands (Bragdon et al., 2011). BMPs belong to the TGF-ß superfamily and have been shown to trigger cell division, differentiation and death, among other cell behaviors (Miyazono, Kamiya, & Morikawa, 2010; R. N. Wang et al., 2014). BMPs are essential regulators of the embryonic development of all metazoans. In fact, loss of BMP signaling invariably results in embryonic lethality (Irish & Gelbart, 1987; Mishina, Suzuki, Ueno, & Behringer, 1995). BMPs are also tightly linked to several human diseases, such as cancer, where they have been described both as oncogenes and tumor suppressors (Bach, Park, & Lee, 2018).

During their biogenesis, pairs of BMP monomers form a disulfide bridge in the endoplasmic reticulum (ER), resulting in a covalently bound dimer. Secreted BMP dimers then act on target cells via the assembly of a heterotetrameric complex comprised of two type I and two type II Serin/Threonine kinase receptors (Little & Mullins, 2009). Recognition of the BMP ligand in the surface of the cell results in type I receptor activation and downstream target gene modulation via phosphorylation of the Smad signal transducer (Mad in *Drosophila*).

*In vivo*, multiple BMP genes are often expressed by the same cell, potentially leading to the formation of different homo- and heterodimers. In most cases, the responding cells can also display different type I and type II receptors. Given that different BMPs have distinct affinities for specific receptors, hetero- and homodimers can result in the formation of unique receptor complexes exhibiting diverse signaling capacities (Little & Mullins, 2009). The multimeric essence of the BMP pathway elements results in a broad combinatorial space, permitting a wide range of signaling outputs depending on the context (Antebi et al., 2017).

Despite their importance, the considerable diversity and redundancy of BMP ligands, coupled with the limitations of current genetic tools, has made it difficult to dissect *in vivo* both the presence and the function of BMP homodimers versus heterodimers.

In contrast to mammals, where more than 20 BMP-encoding genes have been described, the *Drosophila melanogaster* genome possesses three genes: *decapentaplegic* (*dpp*), encoding the vertebrate BMP2/4 ortholog, and *screw* (*scw*) and *glass bottom boat* (*gbb*), encoding BMP5/6/7/8 orthologs. In the embryo, Dpp/Scw heterodimers are thought to be largely responsible for mediating dorsoventral patterning (Shimmi, Umulis, Othmer, & O’Connor, 2005). Indeed, Dpp requires Scw to achieve a normal extracellular distribution (Shimmi, Umulis, et al., 2005; Y.-C. Wang & Ferguson, 2005). Nonetheless, localized expression of Scw or Dpp homodimers in the early embryo are each able to activate signaling at similar levels to the heterodimer (Nguyen, Park, Marqués, & Arora, 1998; Y.-C. Wang & Ferguson, 2005). Direct evidence of heterodimer formation remains missing.

In the larval wing precursor tissue, the wing imaginal disc, both *dpp* and *gbb* have been shown to be involved in patterning and growth (Capdevila & Guerrero, 1994; Khalsa, Yoon, Torres-Schumann, & Wharton, 1998; Zecca, Basler, & Struhl, 1995). In this tissue, *dpp* is eventually expressed in a stripe of cells along the antero-posterior axis. From there, it has been proposed that the protein forms a gradient that instructs concentration-dependent gene expression and ensures tissue growth (Lecuit et al., 1996; Nellen, Burke, Struhl, & Basler, 1996). Loss of *dpp* in the wing disc results in wing patterning defects and significant tissue loss (Barrio & Milán, 2017; Bosch, Ziukaite, Alexandre, Basler, & Vincent, 2017; Spencer, Hoffmann, & Gelbart, 1982; Teleman & Cohen, 2000). In contrast, loss of *gbb* in the wing disc results in patterning defects and mild tissue loss that differ in part from those displayed by the loss of *dpp* (Khalsa et al., 1998), suggesting different but potentially overlapping roles during wing development (Bangi & Wharton, 2006a).

For both *Drosophila* and vertebrates, genetic manipulation of individual BMP genes has been used to address the relative contribution of different dimer types. However, such manipulations change the dosage of the different ligand species relative to one another. Given the ability of different ligand species to signal through distinct receptor complexes, it is possible that changing the relative stoichiometry of different monomers alters the true phenotypic consequences. Moreover, *in situ* localization studies are most often done using ectopically expressed BMP proteins which could alter dimer composition, as well as saturate any regulatory proteins and different receptor complexes.

Here, we combined precise endogenous tagging and direct protein manipulation to explore the distribution, regulation, and function of homo- and heterodimers in the *Drosophila* wing disc. We show that despite their distinct gene expression patterns, both Gbb and Dpp form strikingly similar extracellular gradients. This phenomenon reflects our finding that Gbb depends on Dpp for secretion. Furthermore, our use of extracellular membrane tethering via protein binders reveals that Gbb and Dpp are physically linked in the extracellular space, strongly indicative of heterodimer formation. The same experimental approach failed to detect extracellular homodimers in wild type discs. However, in the absence of endogenous Gbb, Dpp homodimers are detected. Together, our results provide strong evidence that a Dpp/Gbb heterodimer is the prevalent bioactive BMP ligand during wing disc development.

## Results

### Endogenous tagging of BMP ligands reveals coincident extracellular gradients of Dpp and Gbb

To visualize endogenous ligands *in situ*, we used CRISPR/Cas9 editing to introduce small epitope tags into the open reading frame of the endogenous *gbb* locus. Previous attempts tagging BMPs with bulky fluorescent proteins has been shown to result in partial loss of function (Matsuda et al., 2021; Akiyama, Wharton, personal commun.). Hence, we knocked in the short tags HA and OLLAS using ssDonor templates (Fig 1A and material and methods). We also generated a double-tagged *OLLAS:HA:gbb* using the SEED/Harvest technology (Aguilar et al., 2022) (Supp Fig 1A and materials and methods). To visualize Dpp, we made use of a recently generated *HA:dpp* allele (Matsuda et al., 2021) and generated an *OLLAS:dpp* version using the same approach employed originally for HA (see materials and methods). All tags were introduced immediately after the most C-terminal proconvertase cleavage site, in the ligand domain. Adult wings of all the alleles generated did not present any morphological defects in homozygosity (data not shown).

**Figure 1:**
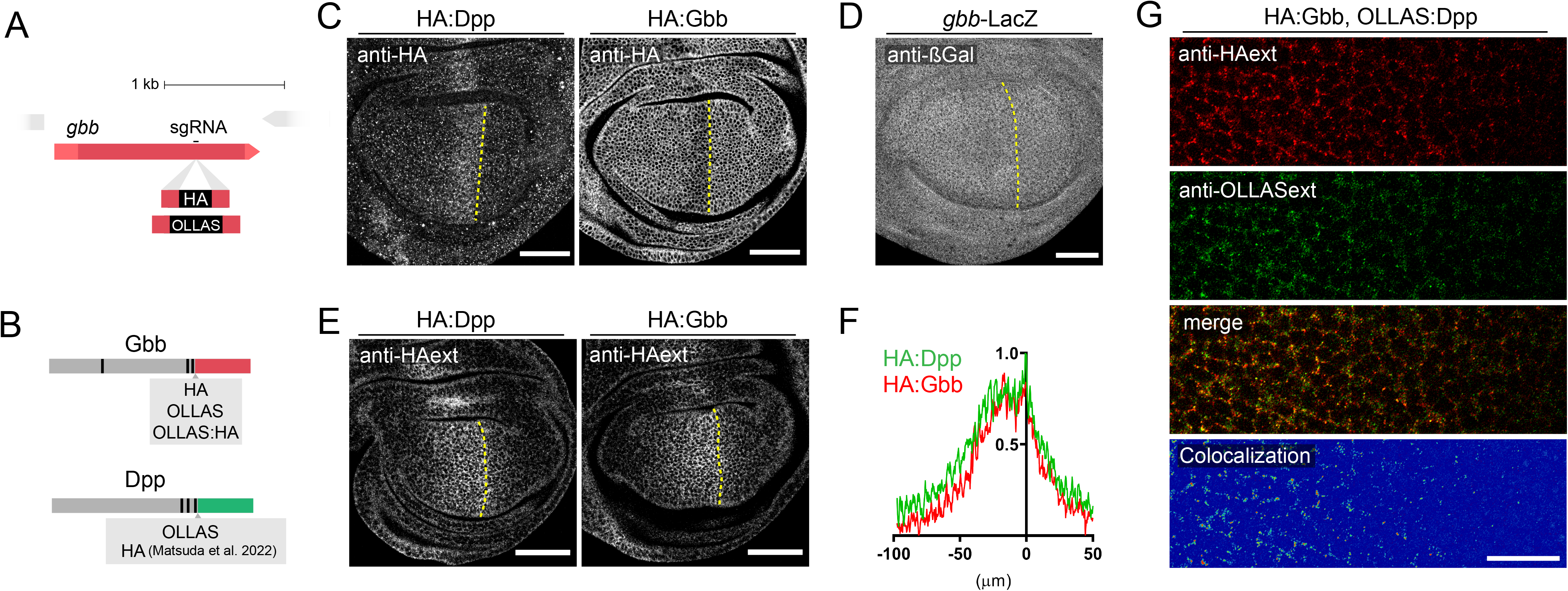
Endogenously tagged Gbb and Dpp form identical extracellular gradients. **A**. Outline of the sgRNA position to manipulate the *gbb* locus via the CRISPR/Cas9 system. **B**. Depiction of the tagging position in Gbb and Dpp. Both genes possess three pro-convertase cleavage sites in their prodomains. All tags were introduced after the last processing site in the beginning of each mature ligand domain (displayed in red for Gbb and green for dpp). **C**. Conventional anti-HA antibody staining of homozygous HA:Dpp (left panel) and HA:Gbb (right panel) wing discs. HA:Dpp is expressed in a narrow stripe of anterior cells along the Antero/Posterior (A/P) compartment boundary. In contrast, HA:Gbb is uniformly detected in the whole wing disc with the exception of a small decrease along the A/P boundary, in the same domain Dpp is expressed. **D**. Anti-ßGal staining of *gbb*-LacZ reporter flies shows the uniform expression of *gbb* in the wing disc. **E**. Extracellular anti-HA antibody stainings in homozygous HA:Dpp and HA:Gbb wing discs. Both HA:Dpp and HA:Gbb form an extracellular gradient, with peak values in the A/P border. **F**. Quantification of the extracellular gradients of HA:Dpp (n=9) and HA:Gbb (n=8) shown in **E**. **G**. Simultaneous extracellular stainings of OLLAS:Dpp (red) and HA:Gbb (green) in the same wing disc. Co-localization of both ligands in the same image (Fake colour scale). Scale bars 50 μm except panel **G** (15 μm).

Consistent with previous reports (Khalsa et al., 1998), immunohistochemical detection of Dpp and Gbb using HA-tagged forms resulted in somewhat complementary patterns in the third instar imaginal wing disc (Fig 1C), HA:Gbb being reduced in the stripe of cells that express HA:Dpp. In contrast to Gbb protein distribution, *gbb* is expressed uniformly as revealed by using a *gbb*-LacZ transcriptional reporter (Fig 1D), confirming previous *in situ* hybridization results (Khalsa et al., 1998). The reduction in Gbb protein compared to *gbb* gene expression levels within the *dpp* expression domain might reflect a post-transcriptional or post-translational process.

Standard immunohistochemistry makes use of permeabilizing agents revealing the localization of both intra- and extracellular proteins. As most of Dpp in third instar wing discs is located intracellularly (Romanova-Michaelides et al., 2022), the presence of low extracellular Dpp levels may be difficult to detect using standard protocols. Eliminating the permeabilization step in standard immunostaining protocols allows for the detection of extracellularly proteins (see Materials and Methods). Using this approach, we found that extracellular HA:Dpp and HA:Gbb exhibit a remarkably similar graded distribution, with maximal levels adjacent to the anterior/posterior boundary (Fig 1 E, F). High-resolution imaging of extracellular OLLAS:Dpp and HA:Gbb showed considerable colocalization of these BMPs in the extracellular space (Fig. 1G). This high degree of extracellular co-localization raised the possibility that the predominant form of secreted BMP ligand is a Dpp-Gbb heterodimer.

### Gbb secretion is dependent on Dpp

BMP proproteins dimerize in the ER of the cells that produce them; therefore, we expect the formation of Dpp-Gbb heterodimers to be limited to those cells in which both genes are expressed. We aimed at determining the source of extracellular Gbb by knocking-down *gbb* expression in either posterior compartment cells by expressing a *gbb*RNAi under the control of the *hh*-Gal4 driver, or in the stripe of cells anterior to the A/P boundary, using *ptc*-Gal4 (Figure 2A). While *gbb* knock-down in the posterior compartment did not alter the graded profile of extracellular HA:Gbb, removing it from the anterior stripe abolished gradient formation (Figure 2A’). Thus, extracellular Gbb is predominately secreted by anterior stripe cells. Given the restricted expression of *dpp* in these cells, we hypothesized that Dpp might be involved in Gbb secretion. We tested this hypothesis by knocking down *dpp* expression in the dorsal compartment with *dppRNAi* driven by *ap*-Gal4 over a period of 18h using a thermo-sensitive Gal80 (longer depletion of *dpp* dramatically altered tissue size and morphology). Dorsal *dpp* knockdown strongly reduced or abolished extracellular Gbb when compared to ventral cells (Fig 2B). This reduction was accompanied by an increase in intracellular Gbb levels (Fig 2B), a buildup reflecting the failure of Gbb to be secreted in the absence of Dpp. The coincident increase of intracellular Gbb and loss of extracellular Gbb was only seen when *dpp*RNAi was expressed in a domain overlapping with the anterior stripe, the domain of endogenous *dpp* expression; no effect was observed when using the posterior driver *hh-*Gal4 (Supp. Fig 2A, B).

**Figure 2:**
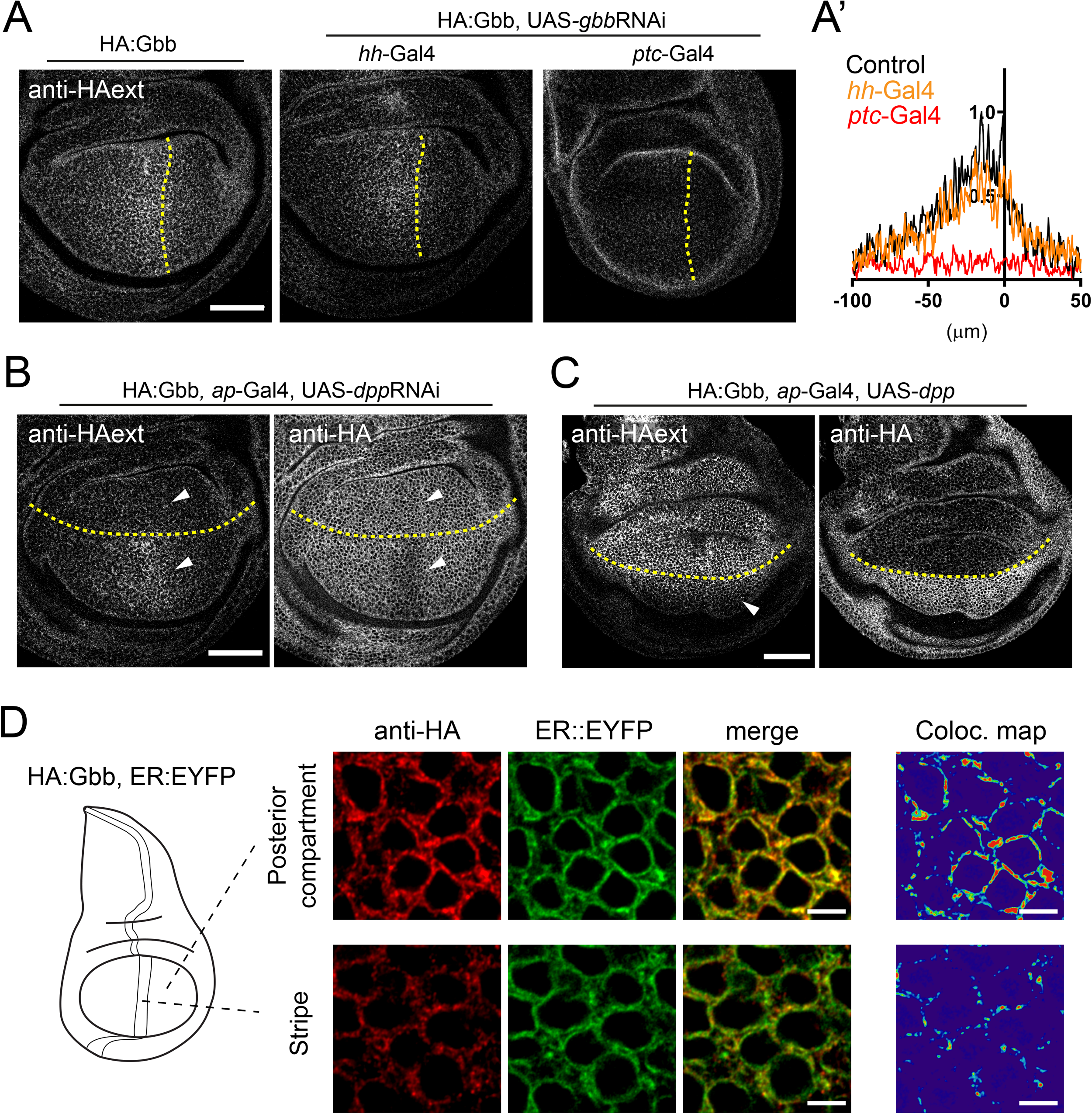
Gbb secretion is dependent on Dpp. **A**. Effect on extracellular HA:Gbb distribution of the expression of *gbb*RNAi in different regions of the wing. Control HA:Gbb staining (left panel). Expression of *gbb*RNAi in the posterior compartment (using *hh*-Gal4) does not influence the extracellular HA:Gbb distribution. Expression of *gbb*RNAi in the A/P stripe, the *dpp* expression domain, (using *ptc*-Gal4) leads to a loss of the extracellular HA:Gbb gradient (mid panel). **A’**. Quantification of the extracellular HA:Gbb gradients with *gbb*RNAi expressed in different domains (control n=4; hh-Gal4 n=4; ptc-Gal4 n=6). **B**. Knock-down of Dpp by *dpp*RNAi in the dorsal compartment leads to a loss of extracellular HA:Gbb when expressed in the ventral compartment using *ap-*Gal4 (right panel, see arrows indicating both compartments). Total anti-HA staining of the same genotype. The reduction of HA:Gbb signal in the A/P stripe is no longer visible in the ventral compartment (left panel, see arrows indicating both compartments). **C**. Overexpression of Dpp in the dorsal compartment leads to massive HA:Gbb secretion from those cells (left panel), invading the ventral compartment (arrow). This leads to a nearly complete depletion of signal in immunostaining of total HA:Gbb in the dorsal compartment, much more pronounced that the reduction in the A/P stripe. Scale bars of panels A, B and C: 50 μm. **D**. Subcellular localization of HA:Gbb in the different compartments and its colocalization with the ER marker *sqh*-EYFP:KDEL. Note that most of Gbb signal in the compartment localizes in ER positive structures around the nucleous. Scale bar 3μm.

We then tested whether the expression of *dpp* was sufficient to trigger ectopic Gbb secretion. Overexpression of *dpp* in the dorsal compartment resulted in a large increase in extracellular secretion of endogenous HA:Gbb from all dorsal cells, creating an ectopic gradient invading the ventral compartment (Fig 2C). In the same genotype, we observed a profound reduction of the intracellular levels of HA:Gbb (Fig 2C). Similar results were obtained when *dpp* was overexpressed in the posterior compartment of the wing disc (Supp. Fig. 2C). The lower level of HA:Gbb seen in the anterior stripe of wild type discs was higher than that seen in cells when *dpp* was overexpressed, suggesting that intracellular Gbb is normally in excess and the level of Dpp determines how much Gbb is secreted. In both *dpp* knockdown and overexpression, *gbb* gene expression remained unaltered (Supp. Fig. 2D).

Consistent with a hypothesis of intracellular ligand retention, HA:Gbb colocalized with an ER marker (Lajeunesse et al., 2004) in both anterior and posterior compartments (Fig 2D). The level of posterior HA:Gbb in ER positive structures was higher than similarly localized HA:Gbb in the anterior stripe (Fig 2D).

### Extracellular probing of BMPs reveals the existence of heteromeric ligand complexes

Collectively, our results are consistent with the hypothesis that Dpp and Gbb form heterodimers, given that both exhibit a similar extracellular distribution and that the presence of Dpp is associated with increased extracellular Gbb. Tools to directly detect the presence of heterodimers *in situ* have been lacking. Therefore, to interrogate the possible direct physical interaction between Dpp and Gbb in the extracellular space, we designed an assay utilizing HATrap, a synthetic membrane-tethered receptor that can bind and trap secreted HA-tagged proteins with high affinity on the cell surface (Matsuda et al., 2021).

We predicted that when expressed in posterior cells (Figure 3A), HATrap would bind and accumulate extracellular HA-tagged peptides at the cell surface (Fig 3B). If the bound extracellular HA-tagged peptides originated from cells of the anterior stripe, they would be trapped and concentrated adjacent to the A/P compartment boundary (Fig 3B).

**Figure 3:**
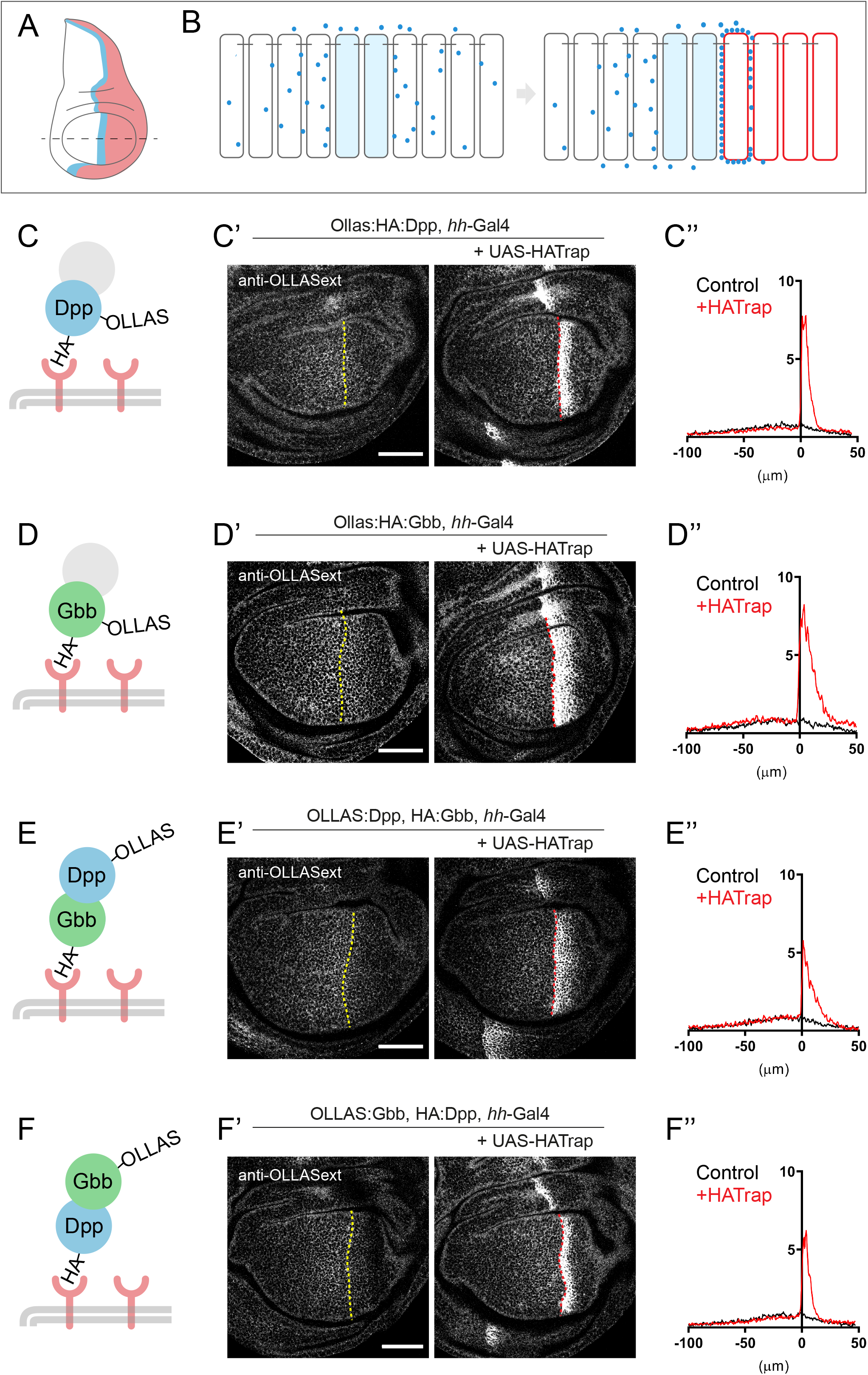
Extracellular tethering of BMP ligands reveals heterodimer existence. **A-B**. Schematic overview of experimental set-up in wing disc. **A**. Gbb and Dpp are both secreted from the anterior stripe of cells along the A/P compartment boundary (blue stripe). Expression of HA-Trap in the posterior compartment (labeled in red) could result in accumulation of stripe derived HA-tagged ligands (for a review on the approach, see (Matsuda, Aguilar, Vigano, & Affolter, 2022)). **B**. In a wildtype tissue, BMP ligands are secreted from source cells and disperse distally, thereby creating an extracellular gradient. Expression of HA-Trap in the posterior compartment leads to accumulation, close to the stripe, of any HA-tagged secreted peptide. **C**. Proof of principle of the system. Expression of HA-Trap with *hh*-Gal4 in an OLLAS:HA:Dpp background. Notice the striking accumulation of Dpp when trap is expressed (right panel) respect no trap being expressed (left panel), as revealed with anti-OLLAS staining. Accumulation occurs in the first few posterior cells adjacent to the A/P border (control n=7; HA-Trap n=7). **D**. Likewise, OLLAS:HA:Gbb can be accumulated upon expression of HATrap by the *hh-*Gal4 driver. In spite of being expressed ubiquitously, OLLAS:HA:Gbb is only accumulated close to the stripe, indicating that only stripe derived peptides are secreted, even if co-expressed with HATrap. Notice the broader accumulation respect panel C (control n=8; HA-Trap n=8). **E**. Trapping of HA:Gbb leads to a inderect accumulation of OLLAS:Dpp in when HATrap is expressed in posterior cells via *hh*-Gal4 (control n=8; HA-Trap n=9). **F**. HA:Dpp trapping in posterior cells leads to an indirect accumulation of OLLAS:Gbb (control n=9; HA-Trap n=9). Scale bar 50 μm.

We first tested the ability of HATrap to accumulate an endogenously expressed, double-tagged OLLAS:HA:Dpp. The use of a double-tagged version is essential for this particular experiment, since the trapping via a given tag prohibits the use of this tag for antibody detection (see: Aguilar et al., 2022). Consistent with previous reports (Matsuda et al., 2021), HATrap expression in posterior cells resulted in a strong accumulation of Dpp just posterior to the A/P boundary, as revealed by anti-OLLAS staining (Fig 3C). Similarly, posteriorly expressed HATrap led to the accumulation of extracellular, double-tagged OLLAS:HA:Gbb, again adjacent to the A/P boundary, strikingly similar to that observed for OLLAS:HA:Dpp, despite the fact that OLLAS:HA:Gbb is expressed throughout the posterior compartment (Fig 3D). As shown in Supp Fig 3A, expression of HATrap masks HA:Gbb when standard immunostaining is performed. One could thus surmise that HATrap could artefactually disrupt the regulated secretion of OLLAS:HA:Gbb when expressed in the same cells; alternatively, the presence of intracellular OLLAS:HA:Gbb could retain HATrap inside the cell, thereby reducing its concentration on the cell surface. The fact that OLLAS extracellular staining was not seen through the entire posterior compartment when OLLAS:HA:Gbb was trapped with HATraping (Fig 3D) indicates that HATrap is not facilitating Gbb secretion. The fact that the band of heightened OLLAS signal is broader than when OLLAS:HA:Dpp is trapped could reflect a reduction in the level of HATrap at the cell surface due to intracellular retention by OLLAS:HA:Gbb.

To test more directly for the presence of heterodimers or, more precisely, for the presence of physically linked Dpp-Gbb molecules, we asked whether trapping of HA:Gbb would lead to an accumulation of OLLAS:Dpp using the set-up described above. HATrap was expressed in posterior cells under the control of *hh-Gal4* in animals with the endogenous *gbb* locus tagged with HA (HA:Gbb) and the endogenous *dpp* locus tagged with OLLAS (OLLAS:Dpp), both in heterozygous conditions. Extracellular immunostaining for OLLAS revealed a band of high signal just posterior to the A/P boundary, showing that trapping of HA:Gbb resulted in an accumulation of OLLAS:Dpp which was produced by the anterior stripe cells (Fig 3E). The reverse experiment, using HA:Dpp and OLLAS:Gbb in the presence of posterior HATrap, produced a similar band of high OLLAS signal. The trapping of HA:Dpp originating from the anterior stripe resulted in the accumulation of OLLAS:Gbb in posterior cells abutting the A/P boundary (Fig 3F). These results provide strong *in situ* evidence that these two BMPs travel together in the extracellular space. The simplest explanation for our observations is that Gbb and Dpp are part of the same complex, forming a Cystein-linked heterodimer that is being trapped. While extracellular heterodimers may be the most plausible explanation, we cannot formally exclude the possibility that our results reflect higher order multimers, or that Gbb homodimers and Dpp homodimers share a common carrier that allows both ligands to be trapped by HATrap.

### BMP homodimers are not detected under physiological conditions

We made use of the same HATrap assay to interrogate the existence of homodimers. To that end, we generated transheterozygous animals that had one *dpp* alele tagged with HA and the other tagged with OLLAS. If homodimers form, the posterior expression of HATrap should result in an accumulation of OLLAS:Dpp (Figure 4A). However, we did not detect an accumulation of OLLAS:Dpp in posterior cells (Fig 4A’ and A’’), showing that under physiological conditions we are unable to detect homodimers. Moreover, this negative result increases confidence that the physical interaction of Dpp and Gbb observed in the experiments described above results from heterodimerization and that higher order multimers are a less likely explanation. We then tested for the presence of Gbb homodimers using the same approach, producing transheterozygotes flies carrying a *HA:gbb* and *OLLAS:gbb* alleles. No extracellular accumulation of OLLAS:Gbb resulted upon HATrap expression in the posterior cells (Supp. Fig. 4A). This is consistent with the model that Gbb secretion from the stripe is induced directly by heterodimerization with Dpp, and not indirectly by the stimulation of homodimer secretion.

**Figure 4:**
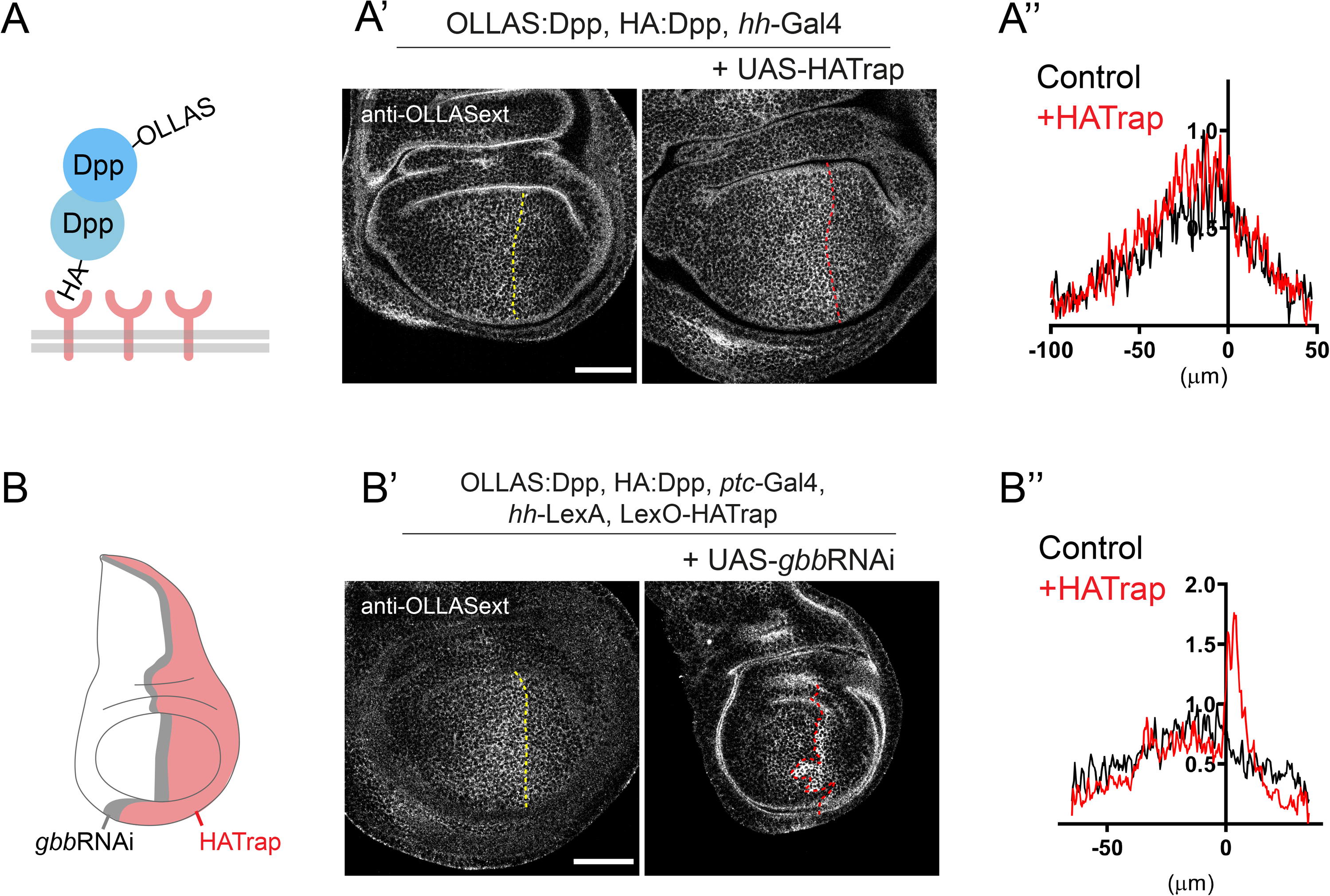
Dpp homodimers can only be detected in the absence of Gbb. **A**. Scheme of the experiment in A’. HAtrap is expressed in Posterior cells using *hh-*Gal4 driver in a background containing both HA:Dpp and OLLAS:Dpp, if homodimers exist HA trapping should lead to OLLAS:Dpp accumulation. A’. Expression of HATrap in posterior cells does not lead to accumulation of OLLAS:Dpp (right panel) compared to control (left panel). **A’’** Quantifications of extracellular OLLAS:Dpp in the absence and presence of HATrap (control n=5; HA-Trap n=7). **B**. Scheme of the experimental setup of B’. HATrap is expressed in the posterior compartment using *hh-*LexA while *gbb*RNAi is expressed in the anterior stripe using *ptc-*Gal4, in a HA-Dpp, OLLAS-Dpp background. **B’**. expression of *gbb*RNAi leads to the indirect accumulation of OLLAS:Dpp when trapping HA:Dpp (right panel) compared to no *gbb*RNAi expression. **B’’**. Quantifications of extracellular OLLAS:Dpp of the panels in B’ (control n=6; HA-Trap n=7). Scale bar 50 μm.

While loss of *dpp* results in dramatic wing defects (Spencer et al., 1982; Teleman & Cohen, 2000), loss of *gbb* has been associated with comparatively minor defects in patterning and growth (Khalsa et al., 1998). This seems to contradict our results, as they suggest that heterodimers are the only bioactive form present in the wing imaginal discs. We hypothesize that the reduced defects in growth displayed by *gbb* mutants are due to compensatory signaling via the Dpp homodimer formed only under such conditions. We tested this hypothesis by trapping Dpp homodimers in the presence and absence of *gbb* expression. To do so, we had to make use of both Gal4/UAS and LexO/LexA expression systems in order to simultaneously express HATrap in posterior cells and *gbb*RNAi in the anterior stripe (Fig. 4B). This setup resulted in the accumulation of OLLAS:Dpp in an *OLLAS:dpp/HA:dpp* background (Fig. 4B’ and B’’), demonstrating that Dpp homodimers are formed in the absence of Gbb, but not in the presence of Gbb (see above).

In order to test whether Gbb homodimers were produced in the absence of Dpp, we trapped HA:Gbb in the presence of OLLAS:Gbb while simultaneously knocking down *dpp* in the anterior stripe (with an 18h expression pulse). In these conditions, OLLAS:Gbb did not accumulate in posterior cells (Supp. Fig. 4B). This result indicates that, in contrast to Dpp, Gbb homodimers are not detected extracellularly in the absence of Dpp.

These results are consistent with our conclusion that, under endogenous physiological conditions, Gbb:Dpp heterodimers are the predominant bioactive ligand in the wing disc. Moreover, the different secretion properties of Dpp and Gbb reconciles our results with the stronger wing phenotypes of *dpp* loss of function compared to *gbb* loss of function.

### Direct manipulation of Gbb results in stronger phenotypes than genetic loss

Our data supports that Gbb/Dpp heterodimer is the main bioactive ligand, Dpp homodimer only being produced upon loss of *gbb*. A direct prediction of this model would be that the genetic loss of *gbb* should produce milder phenotypes than interfering with Gbb at the protein level, once the heterodimers have formed. Direct interference with the heterodimer would not allow for the formation of Dpp homodimers, as happens upon genetic *gbb* loss. HATrap has recently been used to investigate the importance of Dpp dispersal for wing formation (Matsuda et al., 2021). In this case, HATrap was used to trap HA:Dpp on the membrane of Dpp producing cells in an homozygous *HA:dpp* background. This manipulation greatly perturbed signaling in the posterior compartment while the anterior signaling remained largely unaffected (Matsuda et al., 2021).

In the same fashion, we expressed HATrap using the *ptc*-Gal4 driver in an *HA:gbb* homozygous background. This manipulation resulted in strong reduction of the overall phosphorylated Mad (p-MAD) signal, with low levels in the anterior stripe and very little signal in the posterior compartment (Fig 5A). We compared this manipulation with genetic loss by knocking-down *gbb* expression with an RNAi driven by the same *ptc-*Gal4. In this condition, pMad was severely affected with overall reduction in the wing pouch and a narrow peak in the posterior compartment (Fig 5B) as seen when null clones of *gbb* encompass the entire anterior compartment (Bangi 2006). Interestingly, the effect on the posterior pMad is stronger when trapping HA:Gbb than when *gbb* is knocked down (compare Fig 5A and B). Together, these results support the hypothesis that *gbb* genetic loss is rescued by a compensatory mechanism, most likely the ectopic production of Dpp homodimers.

**Figure 5:**
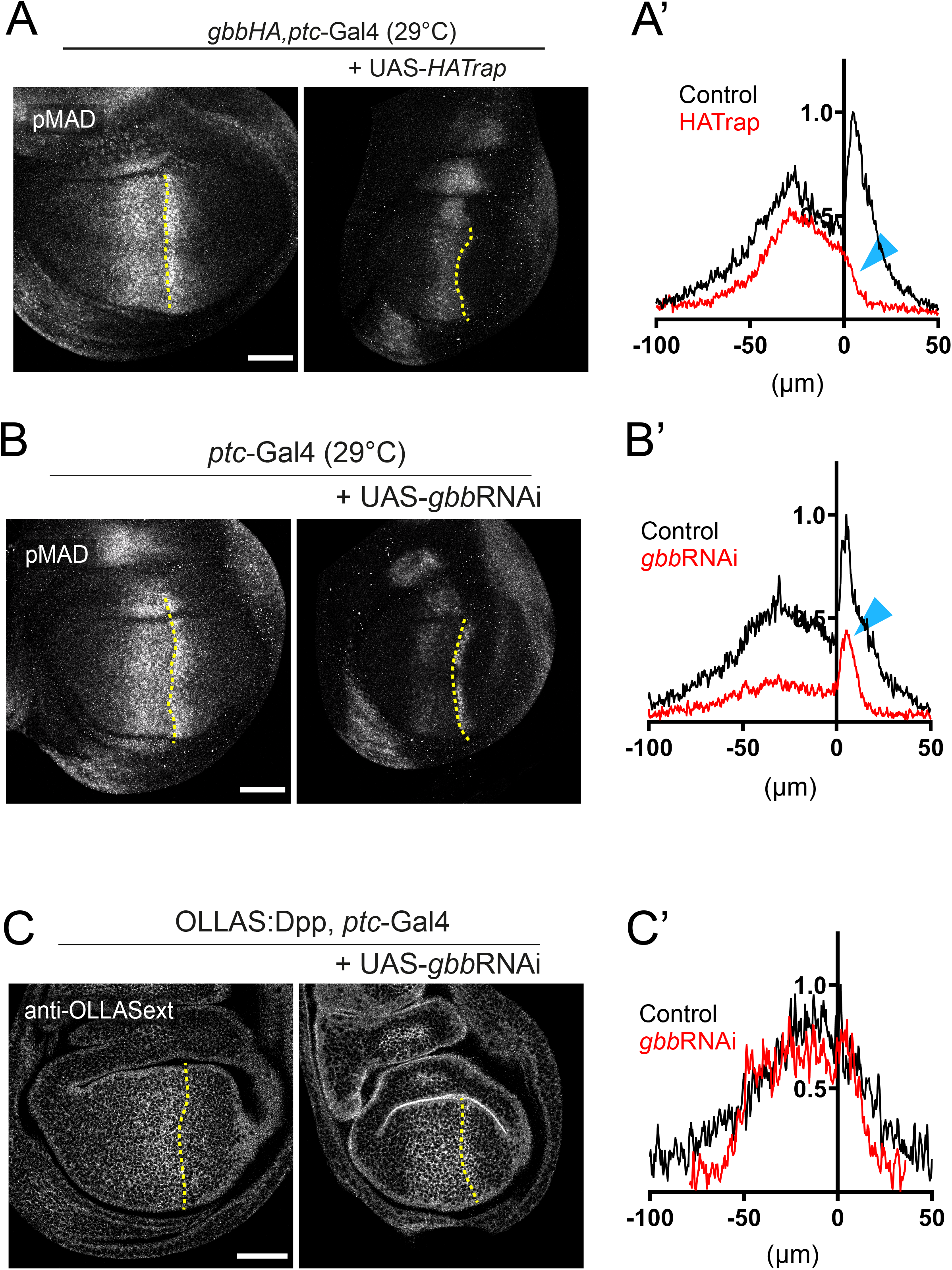
Dpp/Gbb heterodimers are functionally active and ensure long range gradient formation. **A**. Trapping HA:Gbb on the membrane of Dpp producing cells (via *ptc-*Gal4) leads to a strong reduction of p-MAD signaling in the posterior compartment, while the anterior signal is still visible but restricted to the stripe (control n=9; HA-Trap n=13). **B**. Genetic knock-down of *gbb* using *gbb*RNAi driven by *ptc*-Gal4. p-MAD is globaly reduced but the peak in the first cells of the posterior compartment is still visible (control n=7; gbbRNAi n=7). This posterior reduction is especially obvious in the quantification (blue arrows A’ and B’). Experiments in figure A and B were conducted at 29°C. **C**. *gbb* Knock-down in Dpp producing cells (via *ptc-* Gal4) compromises the extracellular OLLAS:Dpp gradient. Leading to a reduction of long-range dispersal of Ollas:Dpp (control n=9; gbbRNAi n=9). Scale bar 50μm.

### Heterodimer formation ensures long range gradient formation

These newly gained insights permitted us to directly compare the heterodimer condition (wild type) versus an exclusive homodimer condition (*gbb* knock-down in the wing pouch, using *gbb*RNAi driven by *nub*-Gal4). We evaluated the extracellular localization of OLLAS:Dpp in both cases in order to assess the distribution of endogenous ligand. Compared to the wild type OLLAS:Dpp extracellular gradient, formed by heterodimers, OLLAS:Dpp homodimers are distributed mainly in the central part of the wing pouch, forming a gradient with abrupt slopes and greatly reduced range (Fig. 5C). The overall anti-HA signal was comparable in both conditions, indicating that OLLAS:Dpp was secreted at similar levels in the absence of Gbb. Similar changes in OLLAS:Dpp distribution were observed when *gbb* was knocked-down in the dorsal compartment (using in the *ap*-Gal4 driver) compared to the ventral compartment (Supp. Fig 5A). Signaling levels and patterning was also severely affected in this condition (Supp. Fig 5B).

### Differential receptor usage of heterodimers and homodimers

Similar to other BMPs, Dpp and Gbb have been proposed to activate signaling through distinct receptor complexes (Bangi & Wharton, 2006a, 2006b; Shimmi, Ralston, Blair, & O’Connor, 2005). However, the relative contribution of each receptor when binding either homo- or heterodimers has never been studied in detail. The wing imaginal disc expresses two genes encoding BMP type I receptors, Saxophone (Sax) and Thickveins (Tkv) and two genes encoding the Type II receptors, Punt (Put) and Wishful thinking (Wit) (Brummel et al., 1994; Childs et al., 1993; Marqués et al., 2002). The active signaling receptor is a heterotetramer composed of two type I and two Type II subunits, therefore multiple receptor combinations are possible in the wing disc. We interrogated the requirement of specific receptors in heterodimer and homodimer conditions (Fig 6A). As described above, the Dpp-Gbb heterodimer is the only detectable form of dimeric ligand present in wild type discs. In turn, *gbb* knock-down results in the exclusive formation of Dpp homodimers. Thus, we knocked-down the different receptors in the presence of Gbb (“heterodimer background”) or its absence (“Dpp homodimer background”) using the wing pouch driver *nub*-Gal4 (Fig 6B). We did not examine Tkv as its requirement for Dpp signal transduction is well established (Ruberte, Marty, Nellen, Affolter, & Basler, 1995; Schwank et al., 2011; Tanimoto, Itoh, ten Dijke, & Tabata, 2000).

**Figure 6:**
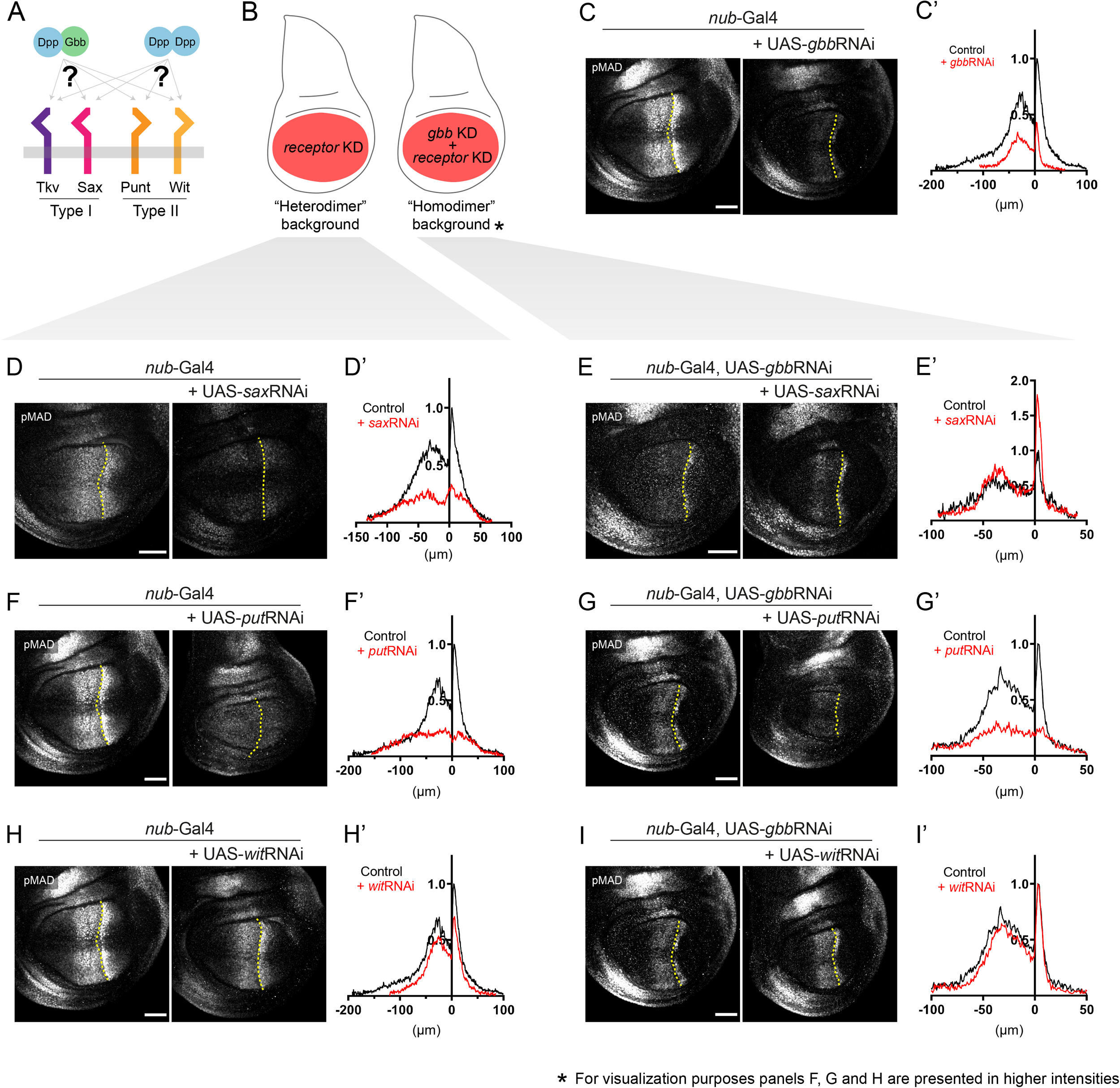
Heterodimer and homodimer use distinct receptors. **A**. Gbb/Dpp heterodimer and Dpp homodimer potentially use different type I and type II receptors in the wing disc. In order to systematically analyze this, we tested the preferential usage of each receptor in a hetero- and homodimer environment. **B**. In a wildtype wing disc, Gbb/Dpp heterodimer are the only form present, while removal of Gbb leads to a Dpp-homodimer environment. In these two conditions, the receptors Saxophone (Sax), Punt (Put) and Wishful thinking (Wish) were knocked-down by RNAi and the resulting pMAD signaling levels compared. **C**. Removal of *gbb* in the wing pouch via *gbb*RNAi with the driver *nub*-Gal4. p-MAD levels and range are reduced in comparison to a wildtype wing disc (control n=9; gbbRNAi n=10). **D**. Knock-down of *sax* by RNAi in a wildtype environment reduced the overall pMAD levels without effecting the low p-MAD levels far from the source (control n=9; saxRNAi n=6). **E**. Removal of *gbb* and *sax* together, did not further reduce p-MAD levels in comparison to *gbb*RNAi alone, but enhanced signaling activity slightly in the posterior compartment, accompanied by a small decrease in range (control n=8; saxRNAi n=7). **F**. Knock-down of *put* in a wildtype wing disc also reduced overall pMAD levels without interfering with the range (control n=9; putRNAi n=8). **G**. Reducing the levels of *gbb* and *put* together lead to a further decrease in pMAD levels, in comparison to the reduction obtained by only *gbb*RNAi (control n=10; putRNAi n=9). **H**. Removal of the receptor *wit* by RNAi in a heterodimer environment only had minor effects on pMAD levels (control n=9; witRNAi n=9). **I**. The simultaneous knock-down of *gbb* and *wit* had little further effect on the already reduced pMAD levels of the *gbb*RNAi background (control n=10; witRNAi n=11). Scale bar 50μm.

Consistent with previous reports (Bangi & Wharton, 2006b), *sax* knock-down strongly affected pMAD amplitude (Fig 6D). Interestingly, this perturbation did not affect the low p-MAD values far from the source, suggesting a minor role in mediating signaling from ligands that presumably act over a long range. In contrast, when both *gbb* and *sax* were knocked-down, pMAD levels where not as reduced as seen in the *gbb* knock-down (compare Fig 6C to Fig 6E). Instead, double *sax*/*gbb* knock-down provoked a mild signal enhancement accompanied by a slight decrease of the range (Fig. 6E). This result shows that in the absence of Sax, Dpp homodimers elicit higher signaling, presumably by binding to receptor complexes containing only Tkv. Knock-down of the Type II receptor *punt* also resulted in a dramatic reduction of p-MAD levels (Fig 6F), as seen by clonal analysis (Letsou et al., 1995; Ruberte et al., 1995), with a change in the shape of the pMad gradient. p-MAD levels are further reduced when both *gbb* and *punt* were knocked-down, significantly flattening the pMad gradient (Fig 6G) compared to that observed in a single *gbb* knockdown (Fig 6C). This confirms that Punt is critical for obtaining normal levels and distribution of BMP signaling in the wing disc (Letsou et al., 1995; Ruberte et al., 1995).

*wit* knock-down only had a minor effect on p-MAD (Fig 6H), suggesting it has a minor role in the transduction of heterodimer signaling in this context, or that its activity can be easily compensated for by Punt. When both *wit* and *gbb* were knocked-down, signal was barely reduced compared to control (Fig 6I). Interestingly, levels in the posterior p-MAD peak were identical in *gbb* single and in *gbb*/*sax* double knock-down, suggesting that Dpp homodimers do not utilize this receptor.

Taken together, our results suggest that in the wild-type wing, BMPs signal through a heterotetrametric complex composed of the type I receptors Tkv and Sax and the type II receptor Punt, with only a minor contribution of Wit. While in the absence of Gbb, Dpp homodimers form receptor complexes composed by Tkv and Punt, with recruitment of Sax having a negative effect on signaling.

## Discussion

BMP signaling is intrinsically combinatorial. At the base of this property is the ability of BMP ligands to form homo- and heterodimers. BMP heterodimers have been shown to be more potent signaling activators than homodimers in several contexts (Hazama, Aono, Ueno, & Fujisawa, 1995; Isaacs et al., 2010; Little & Mullins, 2009; Shimmi, Umulis, et al., 2005; Zhao, Zhao, Koh, Jin, & Franceschi, 2005), probably due to the assembly of distinct receptor complexes (Little & Mullins, 2009). Nevertheless, the different *in vivo* contribution of homo- and heterodimers remains in most cases obscure, mainly due to limitations of classical genetics to dissect *in vivo* dimer composition and/or contribution. In some scenarios, the existence of BMP heterodimers can be inferred by the similar phenotypes caused by the loss of the different ligands (Ray & Wharton, 2001; Shimmi, Ralston, et al., 2005). However, in most cases, both loss and gain of function genetic studies shift the stoichiometry between monomers contributing to the multimeric ligands and thus, fail to reveal the correct bioactive ligand under physiological conditions. In spite of these limitations, a role for heterodimers is suggested by the dominant phenotypes of certain BMP alleles (Ho et al., 2008; Kim, Neugebauer, McKnite, Tilak, & Christian, 2019; Thomas et al., 1997) and by the genetic interactions between different BMP encoding genes (Bangi & Wharton, 2006a; Ray & Wharton, 2001).

Using new reagents coupled with extracellular immunohistochemistry we examined the spatial distribution of Dpp and Gbb in the Drosophila wing primordia and provide evidence that the predominant secreted BMP ligand in the wing imaginal disc is a Dpp/Gbb heterodimer. We propose a model by which the Dpp/Gbb heterodimer biogenesis is regulated (Fig 7A): Gbb is abundant and synthesized in all cells of the wing pouch. Only when co-expressed with Dpp is it no longer retained in the ER, but secreted, as a Dpp/Gbb heterodimer. Dpp, on the contrary, is not retained in the ER and is secreted even in the absence of Gbb. Heterodimerization is favored given the greater relative abundance of Gbb (Fig 2C). The localized expression domain of *dpp* and the regulated secretion of the Dpp/Gbb heterodimer provides a source of bioactive ligand that generates a graded signaling output (Fig 1E) mediated by heterotetrameric receptor complexes containing Sax, Tkv and Punt.

**Figure 7:**
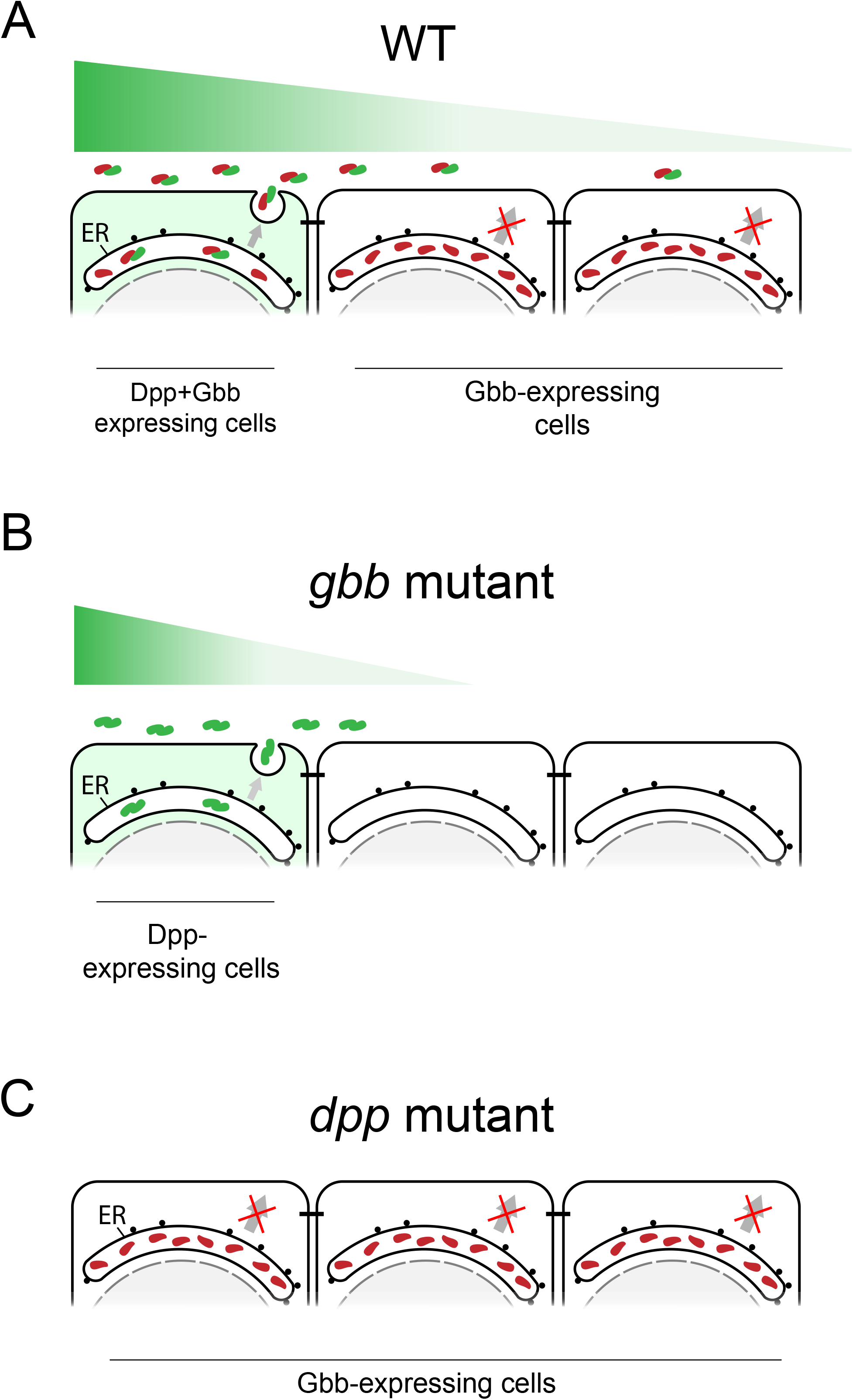
Model of Dpp/Gbb function in the imaginal wing disc. **A**. In physiological conditions, Gbb is synthesized in all cells but retained in the ER. Only in the cells where Dpp is co-expressed (in green on the left), heterodimers will be formed. Heterodimers are the only bioactive form spreading from the stripe, and will result in a long activity gradient. **B**. In absence of Gbb, Dpp will be able to form homodimers. However, these homodimers will have reduced dispersal and signaling capacities. **C**. In absence of Dpp, Gbb will remain retained in the ER, no ligand being secreted, the phenotype will reflect total loss of BMP signal.

Our model can also explain the different phenotypes of *dpp* and *gbb* mutant wings. In *dpp* mutants no bioactive BMP ligands is secreted, as Gbb secretion requires Dpp (Fig 7C). In *gbb* mutants, Dpp can form homodimers and activate signaling, albeit at altered levels and with altered distribution (Fig 4B and Fig 7B). Thus, the phenotype of *gbb* is milder that the phenotype of *dpp*, despite the fact that *in vivo*, all active ligands that we can detect in the with imaginal disc are heterodimers.

Interestingly, mammalian null mutants of BMP2 or 4, the orthologs of Dpp, die during embryogenesis (Winnier, Blessing, Labosky, & Hogan, 1995; Zhang & Bradley, 1996). Conversely, the orthologs of Gbb, BMP5 and 7 have comparatively milder phenotypes, surviving embryogenesis (Dudley, Lyons, & Robertson, 1995; Green, 1958; Kingsley et al., 1992; Luo et al., 1995). This raises the possibility that similar mechanisms to that we describe here are present in other contexts.

### Heterodimerization-dependent secretion

While novel for BMPs, the concept of intracellular retention has been proposed for another member of the TGF-ß superfamily. During early zebrafish embryogenesis, the Activin-like factors Nodal and Vg1 are required for mesoderm induction (Bisgrove, Su, & Yost, 2017; Montague & Schier, 2017; Pelliccia, Jindal, & Burdine, 2017). Vg1 cannot be processed nor secreted in the absence of Nodal (Montague & Schier, 2017). Vg1 is only secreted in the cells where both Vg1 and Nodal are co-expressed. This observations and our model shed light on a conundrum: how can a ubiquitously localized factor, such as Gbb is in the wing imaginal disc (Fig 1B and (Khalsa et al., 1998)) or Vg1 in the early zebrafish embryo (Dohrmann, Kessler, & Melton, 1996; Helde & Grunwald, 1993), confer spatial information to the tissue? The mandatory requirement of heterodimer formation for secretion provides a satisfactory answer. While these factors are essential, it is their localized secretion governed by the formation of a heterodimeric ligand that enables patterning.

The regulated retention of Gbb remains to be explored. Vg1 monomer retention has been proposed to occur mainly through its pro-domain, through exposed cysteines, glycosylated asparagines, and BiP chaperone-binding motifs (Dingal, Carte, Montague, & Schier, 2021). Only when heterodimerization occurs, processing is completed and secretion is initiated.

In the case of Gbb, it has been described that differential processing of the Gbb pro-protein by proconvertases depends on O-glycosylation (Anderson & Wharton, 2017) and at least in the wing disc, the mature ligand retains a large portion of the pro-domain remains attached to the mature ligand (Akiyama, Marqués, & Wharton Kristi, 2012; Anderson & Wharton, 2017). Moreover, cell culture experiments have shown that Gbb can be secreted upon co-expression of Dpp independently of its processing (Anderson & Wharton, 2017). In other contexts, it has been proposed that certain factors can bind BMPs and regulate their secretion (Wilkinson et al., 2003). It could be that a similar factor exists in the wing disc to retain Gbb.

### Signaling range and activity

Loss of *gbb* abrogates long-range BMP signaling in the wing disc (Bangi & Wharton, 2006a; Khalsa et al., 1998). This data led to the postulation that Dpp-Dpp homodimers could mediate short-range signaling, while Gbb-Dpp and/or Gbb-Gbb homodimers mediate long-range signaling (Bangi & Wharton, 2006a). The model presented here argues that the Gbb-Dpp heterodimers are the only bioactive BMP ligand in the wild type wing. Nevertheless, independently on their role under physiological conditions, both studies suggest that Dpp homodimers have limited range of activity. Here, we demonstrate that the distribution of Dpp protein is indeed modified when *gbb* is knocked down (Fig 5C), suggesting that Dpp-Dpp homodimers and Dpp-Gbb heterodimers differ not only in their signaling capacities, but also in their dispersal properties. Differential distribution of homo- and heterodimers has been described in *Drosophila*, both in the early embryo and in the pupal posterior cross vein (PCV) (Matsuda & Shimmi, 2012; Shimmi, Ralston, et al., 2005; Shimmi, Umulis, et al., 2005). In these cases, the differences of distribution have been explained by the existence of associated proteins that impact ligand distribution (Sog/Tsg in the early embryo and Sog/Cv in the PCV) that have different affinities for homo- and heterodimers (Shimmi, Ralston, et al., 2005; Shimmi, Umulis, et al., 2005). Whether such mechanism exist in the wing disc during larval stages remains unexplored.

Another factor that may be responsible for the different dispersal capabilities of homo- and heterodimers is the receptor Tkv. Changes in Tkv levels can alter the range of BMP signaling in the wing (Schwank et al., 2011; Tanimoto et al., 2000).

Indeed, heterozygous *tkv* mutants display a dramatic expansion in the p-Mad gradient (Akiyama et al., 2008). Given the high affinity of Dpp for Tkv, it is possible that the restricted distribution of homodimers reflects Tkv binding compared to that of the heterodimer. In support of this idea, we show that in a Dpp-Dpp homodimer background, loss of the type I receptor Sax, results in a more restricted distribution of p-Mad, perhaps due to increased Dpp/Dpp to Tkv interactions. This result indicates that the Tkv-Dpp interaction is critical in shaping the homodimer gradient.

### Importance of endogenous tagging and protein binders

Classical genetics has been key in revealing critical pathways and untangling biological systems. The power of loss and gain of function studies have permitted the functional dissection of numerous biological mechanisms. Yet, these approaches may fail to detect the contribution of specific players that are part of multimeric complexes and/or compensatory mechanisms. Here, we describe a paradigmatic example of such failings. For years, some assigned Gbb as having a minor role compared to Dpp in wing disc patterning and growth based on its phenotypes and lower sensitivity to changes in gene dosage. Our data presented here demonstrates under endogenous conditions, it is the synergistic action between both Dpp and Gbb that is critical for wild type wing patterning.

In recent years, the advent of CRISPR/Cas9 genome editing has cleared the way for endogenous tagging of genes with unprecedented ease. Tagging not only permits the visualization and biochemical characterization of proteins, but also their direct *in situ* manipulation. To do so, genetically encoded tools based on protein binders have been proposed (Aguilar, Matsuda, Vigano, & Affolter, 2019; Aguilar, Vigano, Affolter, & Matsuda, 2019). These tools permit the acute manipulation of tagged proteins in predictable manners, thus opening the door for experiments that were, thus far, out of reach. With respect to the BMP pathway, the combination of endogenous tagging and protein binders has permitted the interrogation of Dpp dispersal (Harmansa, Hamaratoglu, Affolter, & Caussinus, 2015; Matsuda et al., 2021), the construction of a minimal synthetic morphogen (Stapornwongkul, de Gennes, Cocconi, Salbreux, & Vincent, 2020) and here, the detection of heterodimers. It is by combining these and other emerging tools with the power of genetics that we will better understand how proteins coordinate development.

## Material and methods

### Fly strains

The following fly strains were used in this study: *HA-dpp* (Matsuda et al., 2021), *OLLAS-dpp* (this study), *OLLAS:HA-dpp* (Matsuda et al., 2021*) HA-gbb* (this study), *OLLAS-gbb* (this study), *OLLAS:HA-gbb* (this study), *gbb*^*1*^*/CyO* (Wharton et al., 1999), *gbb*-lacZ (as described in (Akiyama et al., 2012), constructed by replacing the *gbb* coding sequence with lacZ open reading frame), *ptc*-Gal4 (Bloomington stock center (BL)207), *hh*-Gal4, *nub*-Gal4 Gift from Dr Manolo Calleja, *hh*-lexA (Simon et al., 2021), UAS-*gbb*RNAi (BL34898), UAS-*dpp*RNAi (BL33618), UAS-*dpp* (BL1486), *ap*-Gal4 (BL3041), *tubulin*-Gal80ts (BL7016, BL7019, BL7017), UAS/LexAop-HATrap, lexAop-HATrap, UAS-*sax*RNAi (VDRC 46350), UAS-*put*RNAi (BL39025), UAS-*wit*RNAi (BL25949).

### Genotypes by figure

| Figure | Genotype |
| --- | --- |
| Figure 1 C | *yw*; *HA-dpp*/ *HA-dpp*<br>*HA-gbb*/ *HA-gbb* |
| Figure 1 D | *(y)w*; *gbb-lacZ*/+ |
| Figure 1 E | *yw*; *HA-dpp*/ *HA-dpp*<br>*HA-gbb*/ *HA-gbb* |
| Figure 1 G | *OLLAS-dpp*, *HA-gbb*/ *OLLAS-dpp*, *HA-gbb* |
| Figure 2 A | *HA-gbb*/+; UAS-*gbb*RNAi/+<br>*ptc-Gal4*; *HA-gbb*/+; UAS-*gbb*RNAi/+<br>*HA-gbb*/+; *hh-Gal4*/UAS-*gbb*RNAi |
| Figure 2 B | *ap-Gal4*/*HA-gbb*; UAS-*dpp*RNAi/*tub-Gal80<sup>ts</sup>* |
| Figure 2 C | *ap-Gal4*/*HA-gbb*; UAS-*dpp*/*tub-Gal80<sup>ts</sup>* |
| Figure 2 D | *HA-gbb*/+; *sqh-EYFP-ER*/+ |
| Figure 3 C' | *OLLAS-HA-dpp*/+; *hh-Gal4*/+<br>*OLLAS-HA-dpp*/+; *hh-Gal4*/ UAS-lexAop HA-Trap |
| Figure 3 D' | *OLLAS-HA-gbb*/+; *hh-Gal4*/+<br>*OLLAS-HA-gbb*/+; *hh-Gal4*/ UAS-lexAop HA-Trap |
| Figure 3 E' | *OLLAS-dpp*/*HA-gbb*; *hh-Gal4*/+<br>*OLLAS-dpp*/*HA-gbb*; *hh-Gal4*/ UAS-lexAop HA-Trap |
| Figure 3 F' | *HA-dpp*/*OLLAS-gbb*; *hh-Gal4*/+<br>*HA-dpp*/*OLLAS-gbb*; *hh-Gal4*/ UAS-lexAop HA-Trap |
| Figure 4 A' | *HA-dpp*/*OLLAS-dpp*; *hh-Gal4*/+<br>*HA-dpp*/*OLLAS-dpp*; *hh-Gal4*/ UAS-lexAop HA-Trap |
| Figure 4 B' | *HA-dpp*, *ptc-Gal4*/*OLLAS-dpp*; *hh-lexA*, lexO HA-Trap/+<br>*HA-dpp*, *ptc-Gal4*/*OLLAS-dpp*; *hh-lexA*, lexO HA-Trap/UAS-*gbb*RNAi |
| Figure 5 A | *HA-gbb*, *ptc-Gal4*/*HA-gbb*<br>*HA-gbb*, *ptc-Gal4*/*HA-gbb*; UAS-lexAop HA-Trap/+ |
| Figure 5 B | *ptc-Gal4*/+ |
|  | *ptc*-Gal4/+; UAS-*gbb*RNAi/+ |
| Figure 5 C | OLLAS-*dpp*, *ptc*-Gal4/+<br>OLLAS-*dpp*, *ptc*-Gal4/+; UAS-*gbb*RNAi/+ |
| Figure 6 C | *nub*-Gal4/+<br>*nub*-Gal4/+; UAS-*gbb*RNAi/+ |
| Figure 6 D | *nub*-Gal4/+<br>*nub*-Gal4/+; UAS-*sax*RNAi/+ |
| Figure 6 E | *nub*-Gal4/+<br>*nub*-Gal4/+; UAS-*sax*RNAi/ UAS-*gbb*RNAi |
| Figure 6 F | *nub*-Gal4/+<br>*nub*-Gal4/UAS-*put*RNAi |
| Figure 6 G | *nub*-Gal4/+<br>*nub*-Gal4/UAS-*put*RNAi; UAS-*gbb*RNAi/+ |
| Figure 6 H | *nub*-Gal4/+<br>*nub*-Gal4/+; UAS-*wit*RNAi/+ |
| Figure 6 I | *nub*-Gal4/+<br>*nub*-Gal4/+; UAS-*wit*RNAi/UAS-*gbb*RNAi |
| Supplement 2 A | HA-*gbb*/+; *hh*-Gal4, *tub*-Gal80<sup>ts</sup>/+ |
| Supplement 2 B | HA-*gbb*/+; *hh*-Gal4, *tub*-Gal80<sup>ts</sup>/UAS-*dpp*RNAi |
| Supplement 2 C | HA-*gbb*/+; *hh*-Gal4, *tub*-Gal80<sup>ts</sup>/UAS-*dpp* |
| Supplement 2 D | *ap*-Gal4/*tub*-Gal80<sup>ts</sup>; *gbb*-lacZ/UAS-*dpp*RNAi<br>*tub*-Gal80<sup>ts</sup>/+; *ap*-Gal4/+; *gbb*-lacZ/UAS-*dpp* |
| Supplement 4 A‘ | OLLAS-*gbb*/HA-*gbb*; *hh*-Gal4/+<br>OLLAS-*gbb*/HA-*gbb*; *hh*-Gal4/UAS-lexAop HA-Trap |
| Supplement 4 B‘ | *tub*-Gal80<sup>ts</sup>/+; OLLAS-*gbb*/HA-*gbb*, *ptc*-Gal4; *hh*-lexA, lexO HA-Trap/+<br>*tub*-Gal80<sup>ts</sup>/+; OLLAS-*gbb*/HA-*gbb*, *ptc*-Gal4; *hh*-lexA, lexO HA-Trap/UAS-*dpp*RNAi |
| Supplement 4 C‘ | OLLAS-*gbb*/HA-*dpp*, *ptc*-Gal4; *hh*-lexA, lexO HA-Trap/+<br>OLLAS-*gbb*/HA-*dpp*, *ptc*-Gal4; *hh*-lexA, lexO HA-Trap/UAS-*gbb*RNAi |
| Supplement 5 A | Ollas-*dpp*/*ap*-Gal4; UAS-*gbb*RNAi |
| Supplement 5 B | Ollas-*dpp*/*ap*-Gal4; UAS-*gbb*RNAi |

### Generation of different tagged *gbb* and *dpp* alleles

The *gbb* alleles were generated using CRISPR/Cas9. The tagging position for HA/OLLAS/OLLAS-HA knock-ins between the residues 351 and 352 of the *gbb* coding sequence was chosen based on the tagging position described in Anderson & Wharton, 2017. gRNA target sites for this region were identified using the CRISPR Optimal Target finder (http://targetfinder.flycrispr.neuro.brown.edu/). The sequence of the sgRNA targeting the *gbb* locus was: 5’ GCCCAACAACGTGCCGCTGC 3’. The gRNA was cloned by annealing of the following primer pair: 5` TGCAGCCCAACAACGTGCCGCTGC 3’ and 5` AAACGCAGCGGCACGTTGTTGGGC 3’ and restriction ligation into the pCFD5 vector according to (Port & Bullock, 2016). Transgenic flies expressing the gRNA were established by injecting the resulting construct into P(CaryP)attP40 (BL 25709). Efficiency was confirmed by crossing the resulting stock with nub-Gal4, UAS-Cas9. Adult wing of the resulting crossing highly resembled published *gbb* null clones in the wing.

To insert the HA-; and OLLAS-Tag, ssDNA donor templates of 120bps (HA-Tag: 5’-TATGTACAGGGTCTGCATCTGGCAGCTGCGGCCGCCTGCATAGTCCGGGACGTCATAGGGATAGCCGCCCGTGCTCTCCATCGGTTCTAGCAGCGGCACGTTGTTGGGCGACACCGACTTCT-3’; OLLAS-Tag: 5’-TATGTACAGGGTCTGCATCTGGCAGCTGCGGCCGCCTTTACCCATCAGGCGGGGTCCCAGCTCGTTCGCGAAGCCGCTGCCGCCCGTGCTCTCCATCGGTTCTAGCAGCGGCACGTTGTTGGGCGACACCGACTTCT-3’) were ordered from Integrated DNA technologies. In both cases, the PAM sequence of the sgRNA target was changed from GGT to GAT to avoid cutting of the donor template.

To insert the OLLAS:HA-Tag the SEED/Harvest Method (Aguilar et al., 2022) was employed. Briefly, a cassette including 185bp homology arms was synthesized and clone by Genewiz into pUC-GW plasmid. Subsequently, the OLLAS:HA SEED cassette, including the selectable marker was introduced into this vector according to the protocol proposed in: (Aguilar et al., 2022).

nos-Cas9 flies (BL 54591) were crossed with P(*U6-gbb*.gRNA)attP40 males and injected the ssDNA or SEED templates. All the survivors were crossed with *gbb*^*1*^/*CyO*. Out of the following F1 generation, several independent stocks were established with *Xa*/*Cyo*,RFP,*Tb*. Specifically, stocks that became homozygous were genotyped by Single Fly PCR in a subsequent step.

Integration of HA and OLLAS:HA was tested with the following primer pair: 5’-TGCGGCCGCCTGCATAGTCC-3’, 5’-CAAGTGGCTGACCGCC-3’ Integration of Ollas was tested with the following primer pair: 5’-TGCGGCCGCCTTTACCCATC-3’; 5’-CAAGTGGCTGACCGCC-3’ In both cases correct integration resulted in a PCR band of 500bps. SEED stocks were established following the crossing scheme reported in (Aguilar et al 2022).

OLLAS-*dpp* flies were generated as described for HA-*dpp* in Matsuda et al., 2021. Instead of an HA-tag, a fragment encoding for the OLLAS-tag was inserted between the XhoI and NheI sites in the plasmid pBS-attb-Dpp4.4 for injection. The resulting plasmid was injected into *yw* M(*vas*-int.Dm)zh-2A; *dpp*^*MI03752*^/*Cyo*, P23. Transformants were screened and stocks were established again according to Matsuda et al., 2021.

### Immunostainings and image acquisition

For a total antibody staining, third instar larvae were dissected in ice cold PBS (pH 7,2, Gibco™) and immediately fixed for 30 min, at room temperature (RT) in a paraformaldehyde solution (4% PFA in PBS). After fixation, the larvae were extensively washed with PBST (0,3% Triton-X in PBS) to permeabilize the tissue. This was followed by a 1h incubation in blocking solution (5% Normal Goat Serum in PBST) at RT and a consequent primary antibody incubation at 4 °C over-night. Primary antibodies were diluted in blocking solution. The next day, the samples were washed 3× 15 mins with PBST and incubated for 2 hours at RT with the secondary antibody solution. Samples were then washed 3× 15 min with PBST, followed by 3×15min washes with PBS. Fixation, incubation and washing steps were performed with the samples gently rotating.

For extracellular stainings, larvae were dissected in cold S2 media, and incubated with the primary antibody for 1h on ice before fixation. Primary antibodies were diluted in blocking solution (5% NGS in S2 media). To guarantee thoroughly distribution of the antibody, samples were gently mixed by tapping at the tube tube every 10 minutes. Afterwards, the samples were washed at least 10x with PBS, followed by fixation (30min, 4% PFA in PBS). After permeabilization and blocking steps, performed as described above for total staining, either a new total antibody staining was conducted over-night, or the samples were incubated with secondary antibody at RT the same day.

For mounting of imaginal wing discs, samples were transferred to Vectashield, the desired tissues were separated from the rest of the larvae and placed on a glass slide. The samples were then covered with a cover slide and sealed using nail polish. Images were acquired with a Leica SP5 or a Zeiss LSM880 confocal microscope and analyzed using ImageJ.

### Generation of colocalization maps

Images were acquired using the LSM880 confocal microscope in Airscan mode (63x objective, 2.7 zoom). Raw data was then processed using Zen black (Airscan process-Automatic Threshold). Generated .czi files were then opened with ImageJ. Background was subtracted using the BackgroundSubtracter plugin to reduce experimental noise derived from immunostaining. Colocalization map was generated using the Coloc_map.groovy plugin, included in the IMCF_Utilities package.

### Antibodies

The following primary antibodies were used in this study: anti-HA (3F10, Roche, 11867423001; 1:300 for total stainings, 1:20 for extracellular stainings), anti-HA (C29F4, Cell Signaling, 3724; 1:500 for total stainings, 1:20 for extracellular stainings), anti-OLLAS (L2, Novus Biologicals, NBP1-06713, 1:20 for extracellular stainings), anti-phospho-Smad1/5 (41D10, Cell Signaling, 9516; 1:200), anti-Wg (4D4, DSHB, University of Iowa; 1:120), anti-ptc (DSHB, University of Iowa; 1:20), anti-ß-Galactosidase (Abcam AB9361)

The following secondary antibodies were used at 1:500 dilutions in this study: goat anti-rat IgG Fc (FITC) (ab97089, Abcam), goat anti-rabbit IgG (H+L) Alexa Fluor 680 (A-21109; Thermo Fisher), F(ab`)2-goat anti-rabbit IgG (H+L) Alexa Fluor 568 (A-21069; Thermo Fisher), goat anti-rabbit IgG (H+L) Alexa Fluor 488 (A-1108; Thermo Fisher), goat anti-mouse IgG (H+L) Alexa Fluor 568 (A-11004; Thermo Fisher), Alexa Fluor 680 AffiniPure goat anti-mouse IgG, Fcγ fragment specific (115-625-071; Jackson ImmunoResearch), IgY (H+L) Cross-Adsorbed goat anti-chicken, Alexa Fluor 488 (A32931; Invitrogen)

### Quantification of extracellular gradients and pMAD

Of each confocal image, an average projection was created by ImageJ using three sequential slices (Stacks → Z-Project). For each of these average projections a signal intensity profile along the A/P axis was created and collected in Excel. Alignment along the A/P compartment boundary was achieved based on an anti-ptc staining. The average signal intensity profile of each experiment, consisting of intensity profiles of various wing imaginal discs, was created using the script wing_disc-alignment.py (Matsuda et al 2022). Different conditions were then compared using the script wingdisc_comparison.py. The resulting signal intensity profiles were visualized by Prism.

## Acknowledgement

We would like to thank Prof. Alex Schier for comments. The work in the laboratory of M.A. was supported by grants from the Swiss National Science Foundation (310030_192659/1) and by funds from the Kanton Basel-Stadt and Basel-Land. GA and MB were supported by ‘Fellowships for Excellence’ from the International PhD Program in Molecular Life Sciences of the Biozentrum, University of Basel. KAW was supported by NIH R01-GM068118. SM was supported by SNSF Ambizione grant (PZ00P3_180019).

## Figure legends

**Supplement 1: Schematic outline of OLLAS-HA double-tag integration in the *gbb*-locus**. To knock-in the OLLAS-HA double tag into the gbb-locus the SEED/Harvest method described in Aguilar et al., 2022 was used. The gRNA and Cas9 were provided by integrated transgenes. We injected a donor plasmid composed of the following elements: a 3xP3-dsRED selectable marker, to screen the following generation for integration of the SEED cassette, targets for two gRNAs with no cutting sites in the fly genome (1# and 2#), the to be inserted OLLAS-HA tag (split into two parts, with a common repeat) and the homology arms to trigger Homology-dependent recombination (HDR) upon double strand break (DSB) formation. Cas9, together with the *gbb* gRNA will generate a DSB, which will be repaired by HDR using the donor plasmid as a template. Upon insertion of the SEED cassette, the selectable marker is excised through the expression of gRNAs 1# + 2# and Cas9 and the OLLAS-HA are knocked-in into the *gbb* locus

**Supplement 2: Requirement of Dpp for Gbb secretion. A**. Control wing disc displaying the extracellular and total staining for HA:Gbb. **B**. Expression of *dpp*RNAi in the posterior compartment of the wing disc (using *hh*-Gal4) does not influence the extracellular nor total HA:Gbb distribution. **C**.Overexpression of *dpp* in the posterior wing compartment (using *hh*-Gal4) leads to a massive HA:Gbb secretion from those cells. Additionally, a nearly complete depletion of signal for a total HA:Gbb immunostaining can be observed. **D**. Knock-down of *dpp* by *dpp*RNAi or overexpression of *dpp* in the dorsal compartment (using *ap*-Gal4) does not change the expression of the transcriptional reporter *gbb*-lacZ. Scale bar 50μm.

**Supplement 3: HATrap binds and masks the HA epitope of HA:Gbb. A**. Trapping of HA:Gbb in the posterior compartment (using *hh*-Gal4) leads to a masking of the HA epitope in the very same area. Trapping of HA:Gbb does not change the extracellular distribution of OLLAS:Gbb. Scale bar 50μm.

**Supplement 4: The absence of Dpp does not enable the detection of Gbb homodimer.A**. Schematic overview of the experimental set-up in the wing disc. If Gbb homodimer exist in the wing disc trapping HA:Gbb should accumulate OLLAS:Gbb. **A’** Expression of the HATrap in the posterior compartment (using *hh*-Gal4) does not lead to accumulation of OLLAS:Gbb. **A’’** Quantification of extracellular OLLAS:Gbb in the absence and presence of HATrap(control n=9; HA-Trap n=7). **B**. Scheme of the experimental set-up of **B’**. HATrap is expressed in the posterior compartment using *hh*-LexA while *dpp*RNAi is expressed in Dpp producing cells using *ptc-*Gal4, in a HA:Gbb, OLLAS:Gbb background. **B’** Knock-down of *dpp* leads to a loss of extracellular OLLAS:Gbb. B’’ Quantification of extracellular OLLAS:Gbb, while trapping HA:Gbb in the presence and absence of Dpp (control n=3; dppRNAi n=4). **C**. Expression of HATrap in the posterior compartment (using *hh*-lexA) leads to indirect accumulation of OLLAS:Gbb when trapping HA:Dpp (left panel) compared to no accumulation when *gbb*RNAi is expressed in Dpp-producing cells (using *ptc*-Gal4). C’ Quantification of extracellular OLLAS:Gbb, while trapping HA:Dpp in the absence and presence of*gbb*RNAi (control n=9; gbbRNAi n=9). Scale bar 50μm.

**Supplement 5: Knock-down of gbb compromises Dpp signaling activity and long-range gradient formation. A**. Knock-down of *gbb* by *gbb*RNAi in the dorsal compartment of the wing disc (using *ap*-Gal4) leads to a loss of long-range dispersal of OLLAS:Dpp. **B**. Expression of *gbb*RNAi using *ap*-Gal4 reduces pMAD levels drastically in comparison to the ventral control compartment. Scale bar 50μm.

## Notes

### Competing Interest Statement

The authors have declared no competing interest.

