## Supplementary figures and images for "Heterodimerization-dependent secretion of BMPs in *Drosophila*"

### Supplemental Figure 1

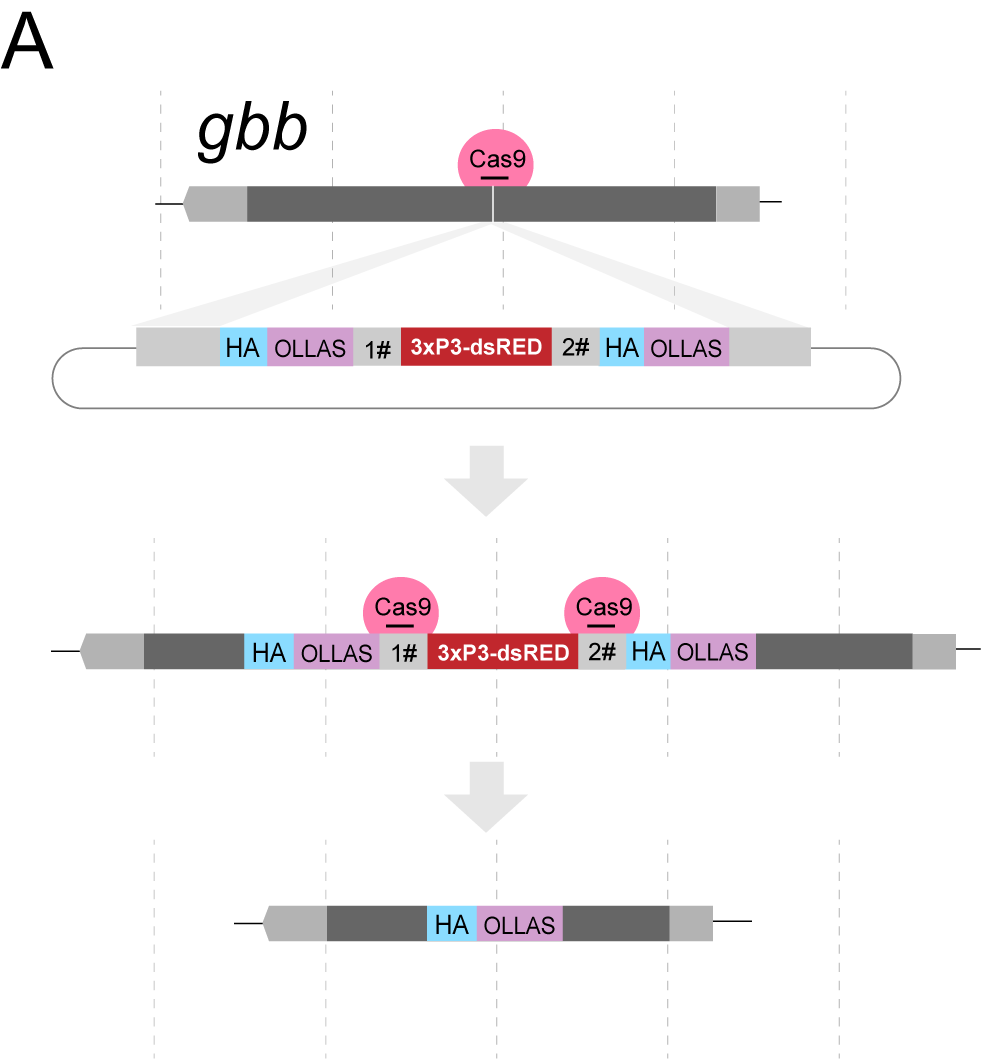

### Supplemental Figure 2

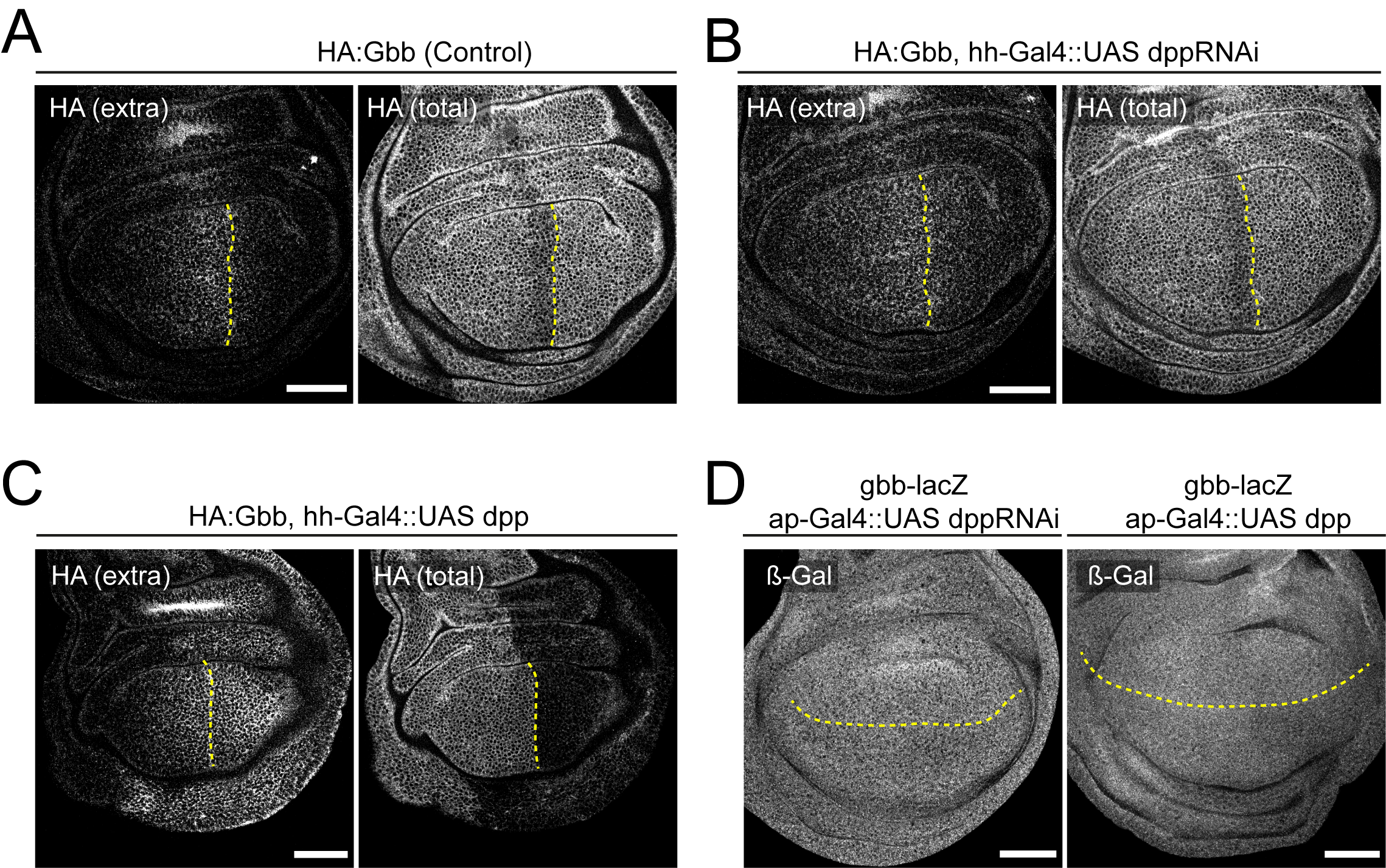

### Supplemental Figure 3

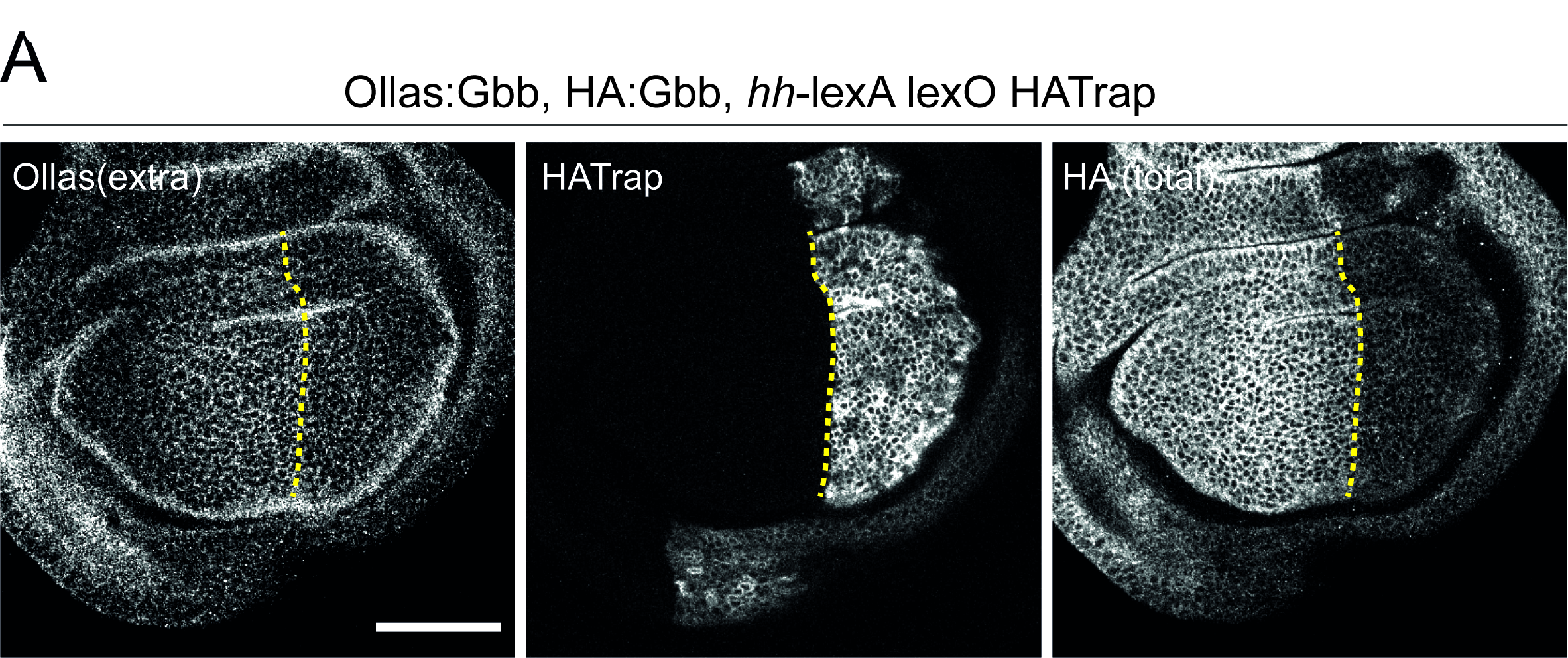

### Supplemental Figure 4

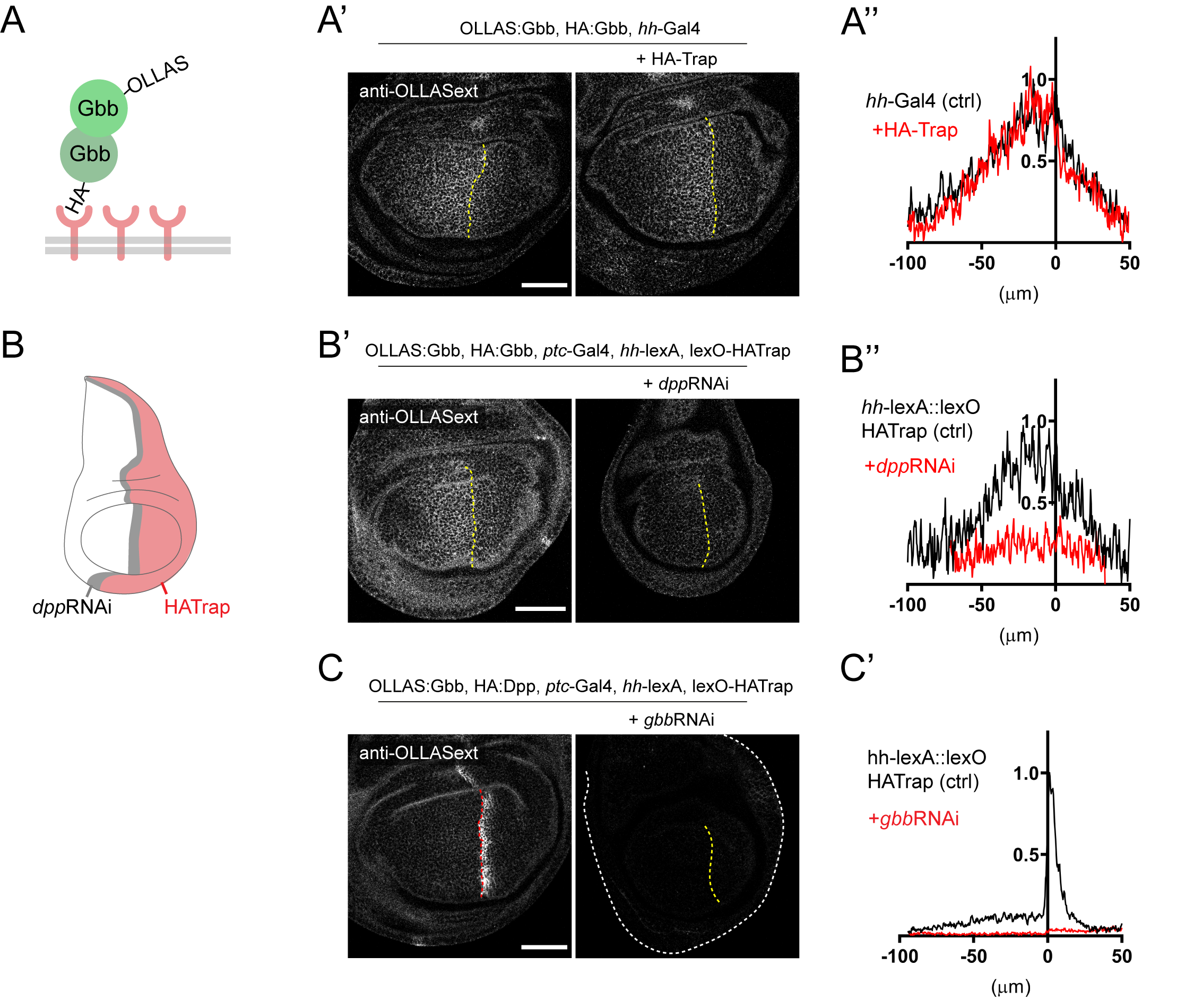

### Supplemental Figure 5

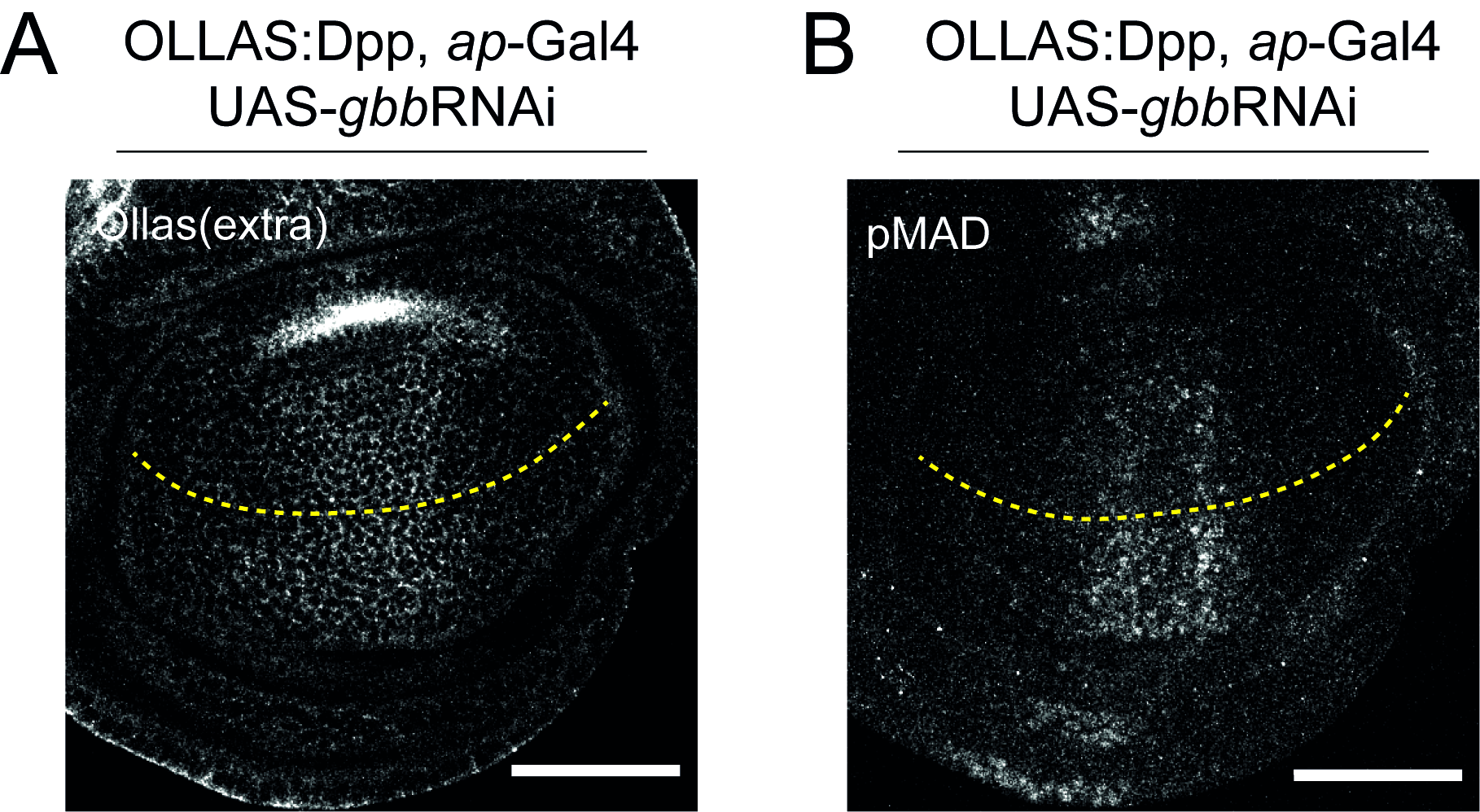
